# TIN2 functions with TPP1/POT1 to stimulate telomerase processivity

**DOI:** 10.1101/435958

**Authors:** Alexandra M. Pike, Margaret A. Strong, John Paul T. Ouyang, Carla J. Connelly, Carol W. Greider

## Abstract

Telomere length maintenance is crucial for cells that divide many times. TIN2 is an important regulator of telomere length, and mutations in *TINF2*, the gene encoding TIN2, cause short telomere syndromes. While the genetics underscore the importance of TIN2, the mechanism through which TIN2 regulates telomere length remains unclear. Here, we characterize the effects of TIN2 on telomerase activity. We identified a new isoform in human cells, TIN2M, that is expressed at similar levels to previously studied TIN2 isoforms. Additionally, we found that all three TIN2 isoforms stimulated telomerase processivity beyond the previously characterized stimulation by TPP1/POT1. Mutations in the TPP1 TEL-patch abrogated this stimulation, implicating TIN2 as a component of the TPP1/POT1 processivity complex. All three TIN2 isoforms localized to telomeres *in vivo* but had distinct effects on telomere length, suggesting they are functionally distinct. These data contrast previous descriptions of TIN2 a simple scaffolding protein, showing that TIN2 isoforms directly regulate telomerase.

## Importance

Telomere length regulation maintains the fine balance between cancer and short telomere syndromes, which are complex degenerative diseases including bone marrow failure and pulmonary fibrosis. The enzyme telomerase maintains telomere equilibrium through highly regulated addition of telomere sequence to chromosome ends. Here, we uncover a previously unknown biochemical role for human shelterin component TIN2 in regulating telomerase enzyme processivity and suggest that TIN2 functions with TPP1/POT1 as a specialized telomeric single-stranded DNA-binding complex. Additionally, CRISPR/Cas9 genome editing revealed a new TIN2 isoform expressed in human cells, and we showed that the three TIN2 isoforms have different effects on telomere length. These findings suggest that previous descriptions of TIN2 as a tethering or bridging protein is incomplete and reveal a previously unappreciated complexity in telomere length regulation. This new perspective on shelterin components regulating telomere length at the molecular level will help advance understanding of clinical manifestations of short telomere syndromes.

## Introduction

Telomere length in human cells is maintained around a tight equilibrium that prevents life-threatening disease. Telomere shortening leads to a characteristic set of degenerative diseases, including pulmonary fibrosis, bone marrow failure, and immune deficiency, collectively called short telomere syndromes(1). In contrast, 90% of human cancers upregulate telomerase, and mutations that increase telomerase levels predispose to cancer(2–4). While we understand many component pathways that regulate telomere length, a detailed integrated mechanism of telomere length regulation is not fully understood.

Human telomeres consist of about 10kb of TTAGGG repeats, that are mostly double-stranded DNA with a single-stranded 3’ overhang, all bound by a protein complex termed shelterin(5). This DNA-protein complex protects chromosome ends, and shelterin both positively and negatively regulates telomere repeat addition by telomerase. The shelterin complex consists of six subunits: two double-stranded DNA binding proteins TRF1 and TRF2(6–9), a single-stranded telomeric binding protein POT1(10, 11), as well as interacting proteins TPP1, TIN2, and RAP1(12–16).

POT1 and TPP1 form a heterodimer that binds single stranded telomeric DNA and stimulates telomerase processivity *in vivo* and *in vitro*(17–19). This stimulation is mediated though the TPP1 OB-fold, which contains conserved TEL-patch and NOB regions that directly interact with the TEN domain of TERT(20–23). Mutations in the TEL-patch abrogate the stimulation of processivity, and compensatory charge swap mutations in TERT restore function(24), suggesting the direct binding of TPP1/POT1 heterodimer to TERT mediates processivity.

TIN2, encoded by the *TINF2* gene, localizes to telomeres through interactions with TRF1, TRF2, and TPP1 (Figure 1A, B). TIN2 interaction with TPP1 is essential for TPP1/POT1 localization and function in cells(25–28). TIN2 also binds to the double stranded DNA binding proteins TRF1 and TRF2(12, 29). Knocking down TIN2 also causes loss of TRF1 and TRF2 at telomeres, suggesting that TIN2 stabilizes TRF1 and TRF2 binding to telomeres(29). Because of its interactions with TRF1, TRF2, and TPP1/POT1, TIN2 has been described as a molecular bridge between the dsDNA- and ssDNA-binding shelterin components. However, it is likely that TIN2 performs additional telomeric functions, as shelterin may consist of distinct functional subcomplexes as implied by genetic and biochemical experiments(29–33).

**Figure 1.**
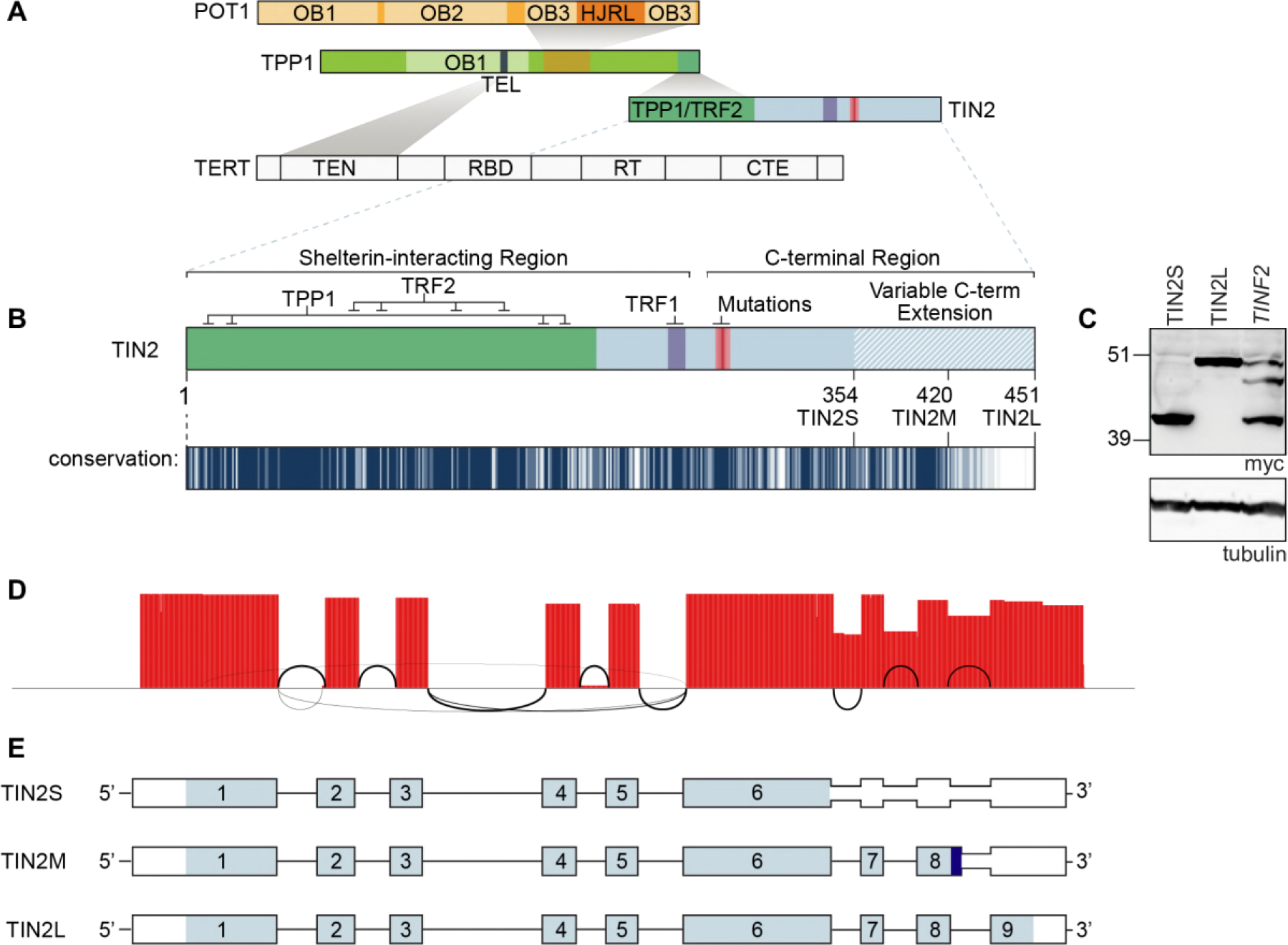
TIN2 has three predominant isoforms in human cells. **a**, Schematic of TIN2, TPP1, and POT1 interaction. The TIN2 N-terminal domain interacts with the C-terminus of TPP1, which is part of a telomerase processivity complex. TPP1 heterodimerizes with the POT1 OB3 domain and also directly interacts with the TERT TEN-domain through a TEL-patch motif. **b**, Detailed schematic of the TIN2 protein. TRF2/TPP1 interaction domain is indicated in green with simplified TPP1 and TRF2 contacts illustrated above. TRF1 FxLxP interaction motif is indicated in purple. The red gradient indicates the patient mutation cluster, where mutated residues cluster but vary in their frequency and disease severity. Light blue hatched region indicates the variable C-terminal extension. Below is a conservation track generated from the values from a multiple sequence alignment with 35 known or predicted TIN2 proteins (see Methods and Supplementary Table 1); colored white = 0, not conserved to navy = 10, highly conserved. **c**, Myc western blot of overexpressed cDNA for TIN2S and TIN2L and the full-length myc-*TINF2* gene. **d**, Sashimi plot of the 3’RACE PacBio sequencing reads aligned to *TINF2*. Height indicates coverage and black lines indicate splicing events, where the line weight corresponds to the frequency of usage. **e**, StringTie-generated TIN2 transcripts from combined data from 293T, HeLa, RPE-1, K562, and LCL cell lines showing TIN2S, TIN2L, and the new isoform, TIN2M. light blue = coding sequence; darker blue = unique TIN2M sequence; white = untranslated region.

*TINF2* mutations cause autosomal dominant inheritance of short telomere syndromes, including dyskeratosis congenita(34, 35) and pulmonary fibrosis(36–38). These mutations are often *de novo*, causing severe disease in patients heterozygous for the mutant allele. These germline missense or nonsense mutations in TIN2 are clustered in a small domain of unknown function (Figure 1B)(34, 35, 39). Within this domain, K280 and R282 are the most commonly mutated residues, and K280E, K280X, R282S, and R282H are the most widely studied TIN2 mutations. These mutant proteins are expressed and stable. In addition to TIN2, mutations in TPP1 and POT1 also cause short telomere disease. The TPP1-ΔK170 mutation deforms the TEN domain binding interface and disrupts the TPP1-telomerase interaction, leading to decreased telomerase processivity(40–42). A POT1-S322L mutation in Coats plus is thought to cause short telomeres through defective telomere replication(43). While mutations in many different genes cause telomere shortening in patients(3), TIN2, TPP1, and POT1 are the only shelterin proteins with mutations identified in short telomere syndromes to date.

Several mechanisms have been proposed for telomere shortening caused by *TINF2* mutations, including defects in telomerase recruitment(44) or decreased telomerase association with the telomere(45). Others have argued for telomerase-independent mechanisms of telomere shortening(46, 47). Several lines of evidence suggest that the TIN2 patient mutations function in a dominant negative manner(34, 37, 45), however, the molecular nature of this effect is not yet understood. To elucidate the mechanism of telomere shortening, we set out to test the biochemical functions of the TIN2 isoforms. We identified a new isoform, TIN2M, and found that all three TIN2 isoforms stimulate telomerase processivity in a TPP1/POT1 dependent manner. The three isoforms had different effects on telomere length when overexpressed in human cells, suggesting functional differences *in vivo*.

## Results

### Identification of a new TIN2 isoform, TIN2M

Human TIN2 is alternatively spliced into two previously described isoforms, TIN2S and TIN2L(12, 48). Most prior work has been performed using the shorter human TIN2S, which was described first, or in the mouse TIN2, which only has one isoform.

TIN2L encompasses all 354 amino acids of TIN2S with an additional 97 C-terminal amino acid residues(48), including a highly conserved domain (Figure 1B)(49). To study TIN2 function *in vivo*, we knocked in an N-terminal myc epitope tag at the endogenous *TINF2* locus in 293T cells using CRISPR/Cas9 genome editing (Supplementary Figure 1A).

Western blots on several edited clones unexpectedly showed three distinct bands, instead of the expected two bands of the known isoforms (Supplementary Figure 1B). To further examine these isoforms, we cloned a myc-tagged full-length *TINF2* gene, including all introns, into an expression vector with the CMV promoter. In transfected cells overexpressing this construct alongside TIN2S or TIN2L cDNA, we again observed an intermediate sized band at approximately 47 kDa. (Figure 1C).

To test whether this band corresponds to an alternatively spliced TIN2 isoform, we used a modified 3’ RACE with PacBio sequencing to identify all full-length expressed isoforms in human and mouse cells. In 293T cells, TIN2S and TIN2L cDNAs were identified along with a third major isoform, which would encode the expected molecular weight for the unknown protein. We termed this isoform TIN2M for TIN2 “medium”. TIN2M results from retention of the last intron, between exons 8 and 9, that encodes 13 amino acids of unique sequence (_408_-VSGKEQKAGKGDG-_420_) before reaching a stop codon (Figure 1D-E). Sequence read counts indicated that TIN2M and TIN2L mRNAs were expressed at similar levels, while TIN2S had 2-fold greater representation than each of the others, suggesting it is the predominant transcript (Figure 1D and Supplementary Figure 2A). 3’ RACE further showed that TIN2M was present at similar levels in four other human cell lines (HeLa, K562, RPE-1, and a newly derived LCL) (Figure 1E). In addition to these three major isoforms in human cells, we identified a number of additional recurrent exon skipping, intron retention, and alternative polyadenylation site usage events, including exon 2 skipping described previously(50) (Supplementary Figure 2A). We found that two different mouse strains (C57BL/6 and CAST/EiJ) expressed just one TIN2 isoform that is most similar to TIN2L, as previously described(48, 51) (Supplementary Figure 2B).

**Figure 2.**
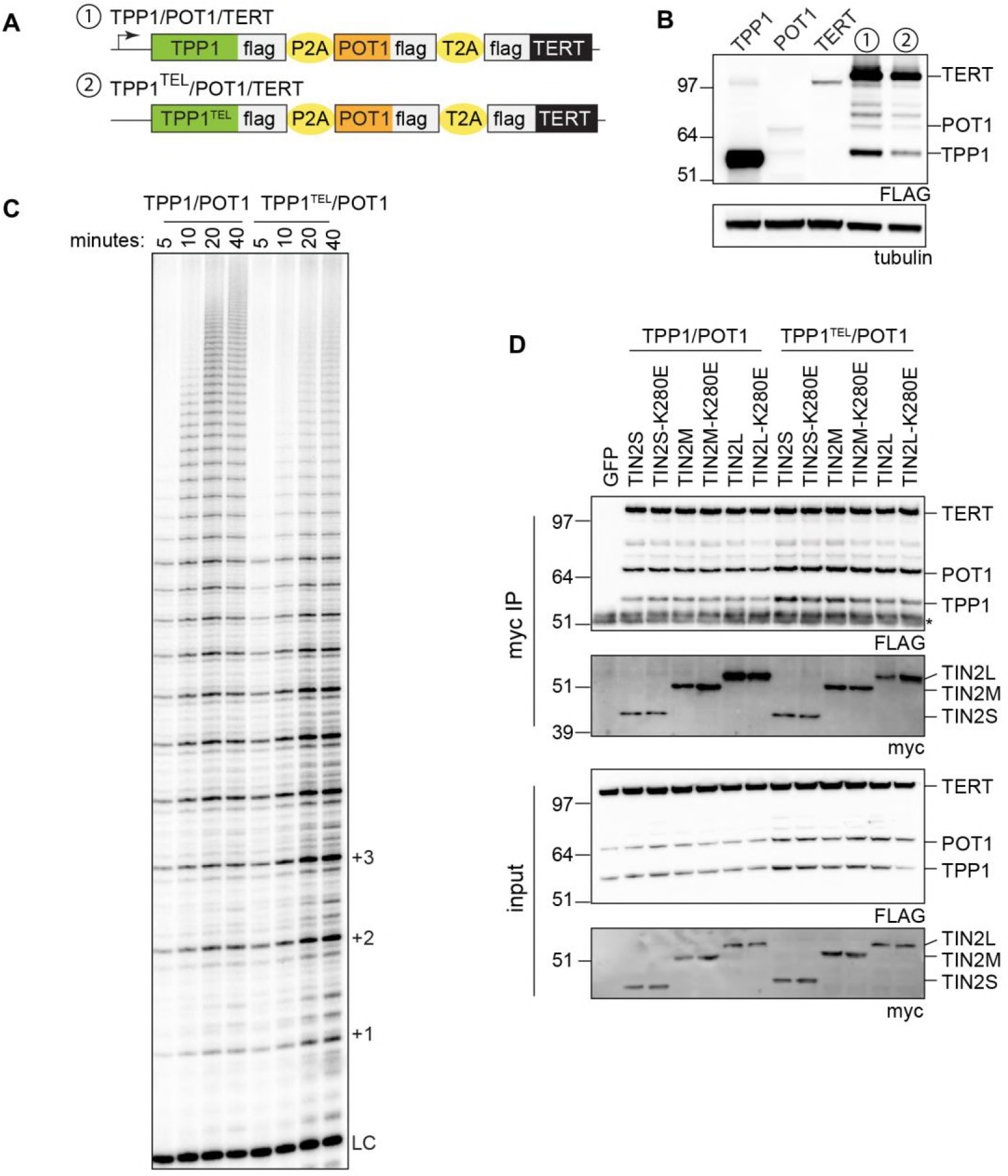
TIN2 isoforms form a complex with TPP1/POT1 that binds telomerase and is not disrupted by the K280E mutation. **a**, Expression cassettes used in this study. All cassettes are expressed by the CMV promoter in the pcDNA5/FRT backbone. Telomerase assay cell lines were generated as described in the Methods. **b**, Western blot of individually transfected TPP1, POT1, and TERT cDNAs next to telomerase assay cell lines numbered as in **a**. FLAG bands above POT1 are unidentified but may be TERT degradation products. **c**, Telomerase assays stopped at 5, 10, 20, and 40 minutes for each cell line. Telomere repeats are indicated by +1, +2, etc. LC = loading and purification control. **d**, Co-immunoprecipitation of TERT, TPP1, and POT1 with TIN2 using anti-myc agarose beads in both telomerase assay cell lines 1 and 2 transfected with TIN2S, TIN2M, or TIN2L. Similar co-IP levels were observed in both WT and mutant constructs. *, IgG bands.

Evidence for expression of TIN2M was also found in publicly available data from PacBio IsoSeq of MCF-7 breast cancer cells (http://www.pacb.com/blog/data-release-human-mcf-7-transcriptome/). Additionally, genome-wide ribosome profiling data from GWIPS-viz shows ribosome peaks present in the unique coding region of the TIN2M retained intron(52). TIN2M and TIN2L, but not TIN2S, contain the recently identified CK2 phosphorylation site(49). All three of the expressed isoforms contain the documented cluster of telomere syndrome patient mutations and the other known interaction domains, suggesting that any of these three isoforms could mediate the short telomere phenotypes seen *in vivo*.

### TIN2 cooperates with TPP1/POT1 to stimulate telomerase processivity

TIN2 interacts directly with TPP1, a processivity factor that heterodimerizes with POT1 and directly binds telomerase through the TPP1 TEL-patch domain(18, 20–22, 53). To examine whether TIN2 affects telomerase activity or processivity, we adapted the cell-based system overexpressing TERT, TR, POT1, and TPP1 used by Nandakumar *et al* (20). By co-overexpressing TERT, TR, TPP1, and POT1 in cells, cell lysates can be used in direct telomerase activity assays(18–20, 54). Because endogenous telomere proteins are expressed at low levels, the telomerase activity observed in this system result from the overexpressed proteins. We adapted this system to generate cells constitutively expressing TERT, TR, TPP1, and POT1, where TIN2 can be introduced by transient transfection.

For reproducible overexpression of the protein components, we created a polycistronic expression cassette containing FLAG-TPP1, FLAG-POT1, and FLAG-TERT separated by 2A peptides (Figure 2A and Supplementary Figure 3A). As a negative control, we mutated the TPP1 TEL-patch (TPP1 E169A/E171A)(20), referred to here as TPP1^TEL^, to test whether any effects of TIN2 are mediated through TPP1/POT1 stimulation of telomerase (Figure 2A and Supplementary Figure 3B). Then, we generated a clonal cell line overexpressing TR in 293TREx FLP-in cells, into which we integrated the respective expression cassette at a unique genomic locus using the FLP-in system. The resulting cell lines are referred to as TPP1/POT1/TERT and TPP1^TEL^/POT1/TERT (Figure 2A-C).

**Figure 3.**
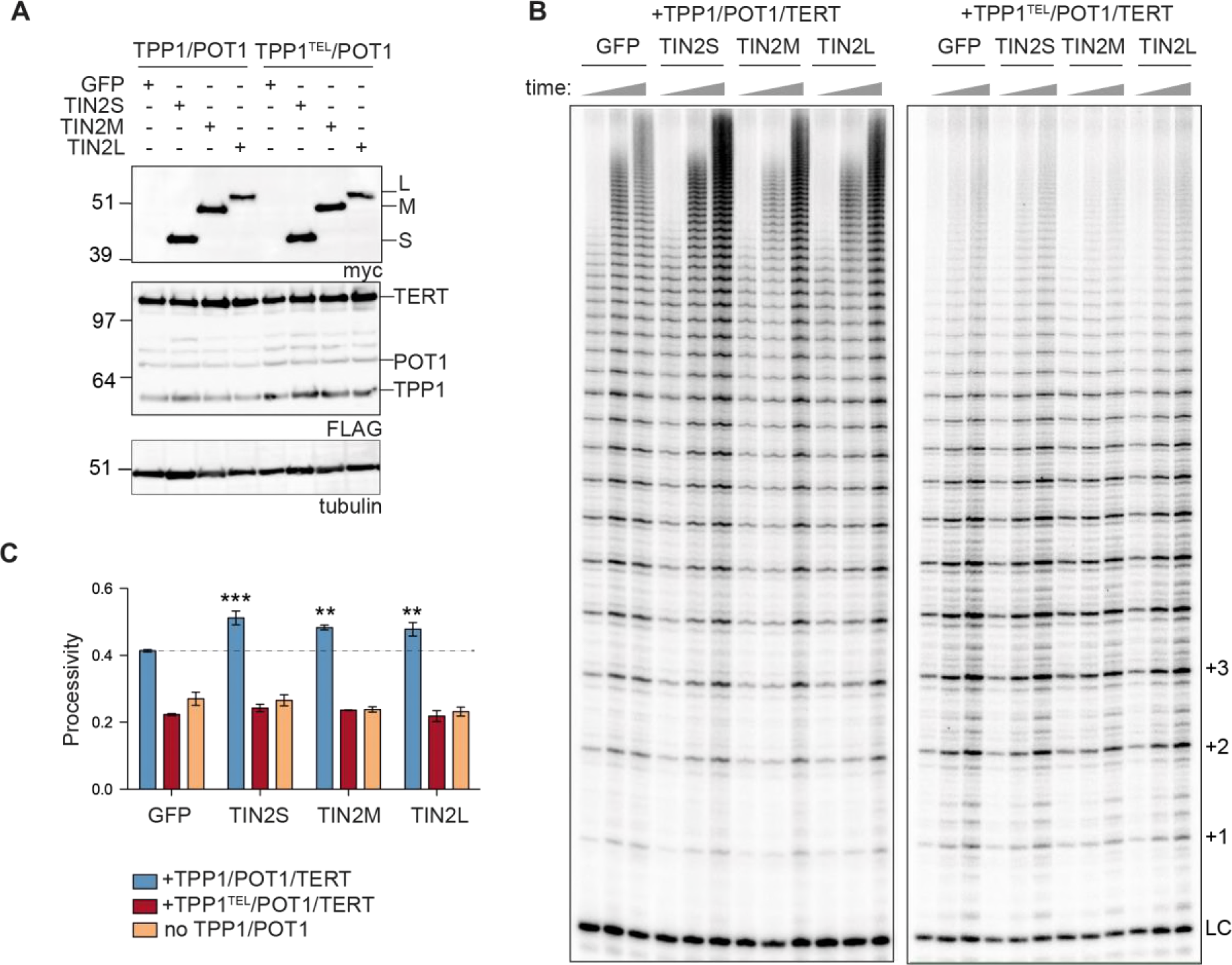
TIN2 stimulates telomerase processivity beyond the TPP1/POT1 stimulation. **a**, Western blots of GFP and myc-TIN2 isoform transfections into TPP1/POT1/TERT (left) or TPP1^TEL^/POT1/TERT (right) cell lines. FLAG bands above POT1 are unidentified but may be TERT degradation products. **b**, Telomerase assays stopped at 10, 20, and 40 minutes. Quantification is shown in (C). Increasing time indicated by the grey triangle. LC, loading and purification control; +1, +2, +3 indicates repeat number. **c**, Mean processivity values from 3 independent telomerase assays at the 40 minute timepoint using the 15+ processivity method (see Methods). Orange bars are from a cell line overexpressing TERT/TR but not TPP1/POT1 (Supplementary Figure 6). Data was analyzed with a one-way ANOVA and Bonferroni’s Multiple Comparisons test against the GFP control. n=3 independent transfections per cell line indicated. Error bars represent SD. **, p<0.01; ***, p<0.001.

To examine the interaction of TIN2 with TPP1/POT1 and telomerase, the three TIN2 isoforms were individually transfected into each cell line. Each of the three TIN2 isoforms reproducibly co-immunoprecipitated with TPP1/POT1 and TERT in reciprocal pull downs of either myc-TIN2 or FLAG-TPP1/POT1/TERT (Figure 2D and Supplementary Figure 4). We observed no change in co-immunoprecipitation of TPP1/POT1 and TERT with TIN2 when any of the three isoforms carried one of the common patient mutations, K280E (Figure 2D), as previously reported for the TIN2S isoform(44, 55). Telomerase activity can be detected in these co-immunoprecipitations, suggesting that the telomerase in complex with TIN2 is active (data not shown). We conclude that all three isoforms of TIN2 are interacting with TPP1/POT1 in complex with active telomerase, and the K280E patient mutation does not disrupt this interaction.

**Figure 4.**
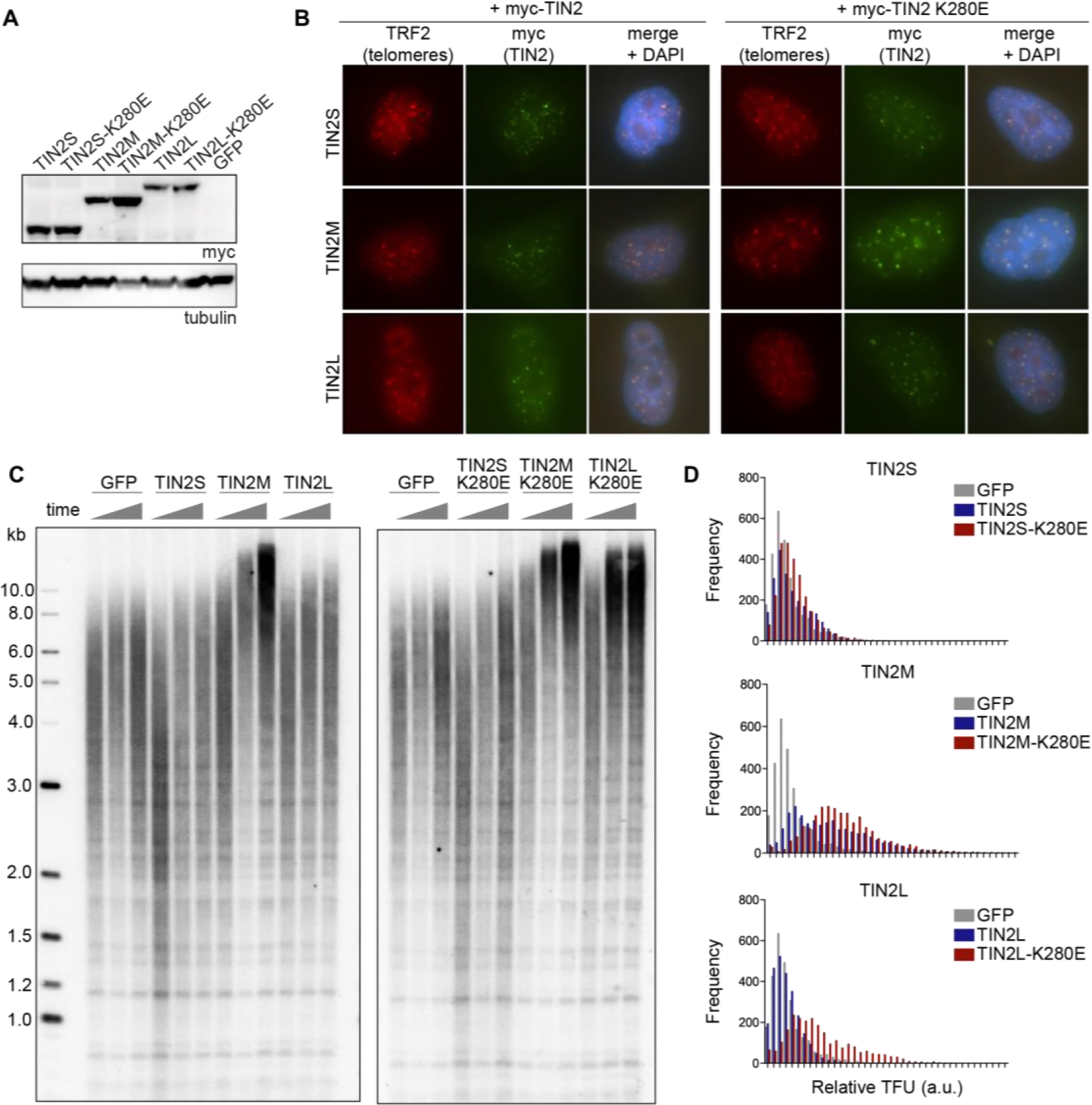
TIN2 isoforms localize to telomeres but have different effects on telomere length. **a**, Western blot of myc-TIN2 overexpressing HeLa FLP-in cell lines. **b**, Immunofluorescence of TIN2 expressing cell lines. TRF2 marks telomeres (red), anti-myc antibody marks TIN2 (green), and nuclei were counterstained with DAPI. Merge image shows telomeric foci with colocalized TRF2 and TIN2 staining. **c**, Telomere Southern blot of genomic DNA from HeLa-TIN2 cell lines. Three timepoints indicated by grey triangles refer cells harvested at 3, 8, and 13 weeks in culture. Left, 2-log ladder values in kb. **d**, Histograms of telomere intensities from quantitative telomere FISH (q-FISH) on late passage metaphase chromosomes in the indicated cell lines, separated by isoform. The same GFP sample is plotted on each graph. Relative telomere fluorescence units (TFU) on the x-axis reflects telomere length. In each, grey = GFP, blue = TIN2-WT, red = TIN2-K280E.

To examine the effects of TIN2 on telomerase activity, we transfected the myc-tagged full length gene, myc*-TINF2*, into the TPP1/POT1/TERT cell line. All three isoforms were expressed from the myc-*TINF2* construct and the lysates showed an increase in processivity compared to the GFP control (Supplementary Figure 5), suggesting that TIN2 enhances telomerase processivity over the effects of TPP1/POT1 alone. To determine if specific TIN2 isoforms are required for this stimulation, we independently transfected each isoform into the TPP1/POT1/TERT cell line (Figure 3A). We found a reproducible 10-20% stimulation of telomerase processivity with each of the three N-terminally tagged isoforms (Figure 3B-C). To test whether TIN2 stimulation of telomerase depends on TPP1/POT1, we transfected TIN2 into TPP1^TEL^/POT1/TERT cells and separately into a TERT-only cell line overexpressing TERT/TR but not TPP1/POT1. We found no stimulation of telomerase processivity in either of these cell lines (Figure 3A-C and Supplementary Figure 6), suggesting the stimulation is dependent on TPP1/POT1. Our results indicate that TIN2 cooperates with TPP1/POT1 to stimulate telomerase processivity.

**Figure 5.**
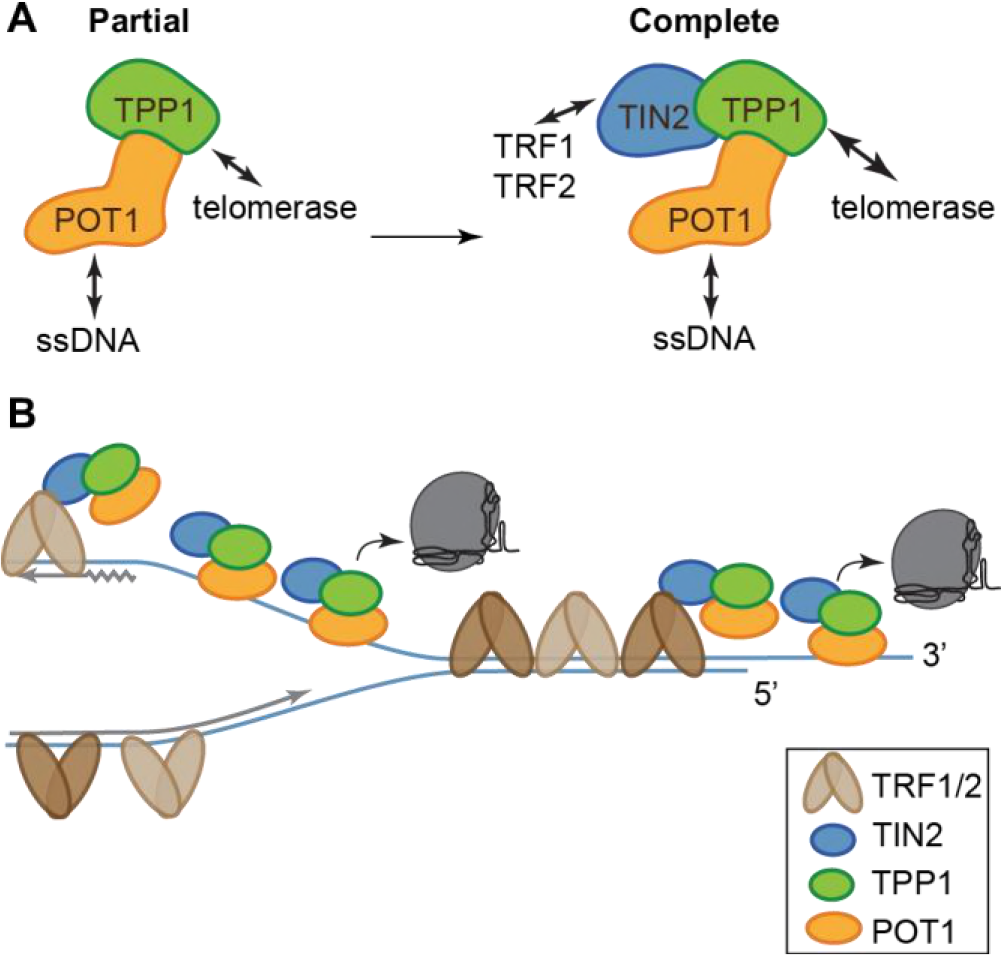
TIN2/TPP1/POT1 is a stable shelterin subcomplex. **a**, TIN2 completes the telomerase processivity complex. TIN2 enhances TPP1/POT1 stimulation of telomerase, forming a heterotrimeric processivity complex that is recruited to the telomere through TRF1/2 interactions. **b**, A dynamic, heterogeneous distribution of shelterin proteins across the length of human telomeres coordinates telomere length maintenance. TRF1 and TRF2 may direct TIN2/TPP1/POT1 to single-stranded DNA both at the telomere overhang and within the replication fork, aiding its roles in fork progression and telomerase stimulation.

Because all three TIN2 isoforms stimulated telomerase to the same extent in a TPP1/POT1 dependent manner, we tested whether patient mutations TIN2-K280E, TIN2-R282S, TIN2-R282H, or TIN2-K280X affect processivity. In some instances, we found that TIN2 mutants were deficient at stimulating telomerase activity, but this result was variable both in whole-cell lysates and in TIN2 co-immunoprecipitations (Supplementary Figure 5 and data not shown). Because the mutants are dominant-negative *in vivo*, we tried co-expressing wild-type TIN2 with a mutant TIN2, but there was no change in processivity stimulation in this setting (Supplementary Figure 7). Although the patient mutations did not reproducibly affect telomerase processivity, we have identified a previously unknown role of TIN2 isoforms in telomerase processivity stimulation that changes the understanding of TIN2’s role in telomere length regulation.

### TIN2 isoforms localize to telomeres and have different effects on telomere length

Since all TIN2 isoforms stimulated telomerase in a TPP1/POT1 dependent manner, we examined whether they function differently in human cells. TIN2S and TIN2L have been demonstrated to localize to telomeres *in vivo* through interaction with TRF1 and TRF2(12, 48). Patient mutations did not disrupt the localization of TIN2S(44), but localization of TIN2L with patient mutations has not been reported. To determine whether TIN2M localizes to telomeres and whether patient mutations affect localization of TIN2M or TIN2L, we examined the localization of each isoform with or without the K280E patient mutation.

Because of the alternative splicing of *TINF2* transcripts, we could not test expression of individual isoforms at the endogenous locus. Instead, we stably overexpressed cDNA encoding TIN2S, TIN2M, or TIN2L with or without the K280E patient mutation in HeLa-FRT Flp-in cells (Figure 4A). Using this system, the expression constructs were integrated at a unique genomic locus, and isogenic, polyclonal cell lines were selected. Western blot analysis showed similar expression levels of TIN2S, TIN2M, and TIN2L that was not affected by the K280E mutation (Figure 4A). Using indirect immunofluorescence, we found that all three isoforms, with or without the K280E patient mutation, showed discrete foci that co-localized with TRF2, indicating that they each localize to telomeres *in vivo* (Figure 4B).

Previous work has shown that overexpression of wild-type TIN2S had little effect on telomere length, while overexpression of TIN2S-K280E, TIN2S-R282S, or TIN2S-R282H decreased telomere length(12, 44). A recent study showed overexpression of TIN2L resulted in some increase in telomere length(49). Having these isoform-specific polyclonal TIN2 overexpressing cell lines in hand, we examined how the TIN2 constructs affect telomere length. We passaged these cells and monitored telomere length by Southern blot and q-FISH analysis. TIN2S, TIN2S-K280E, and the control GFP cell lines showed no significant changes in telomere length over time (Figure 4C-D). In contrast, TIN2L showed some telomere elongation, and TIN2M, TIN2M-K280E, and TIN2L-K280E showed significant increases in telomere length (Figure 4C-D). The excessive telomere elongation resembles the telomere elongation in a number of TPP1/POT1 loss of function mutants(12, 13, 31, 44, 56). These telomere length changes are not due to clonal variation, as our cells are a mixed clonal population of isogenic cells. Telomere length effects appear to differ between immortalized cell lines, as TIN2 overexpression had little effect on telomere length in 293TREx cells (data not shown). The observation that the different TIN2 isoforms and mutants do have strong yet different effects in HeLa cells suggests that these isoforms play different functional roles in the cell.

We next examined telomere aberrations in blinded q-FISH images. We saw no changes in signal-free ends, PQ ratios, sister telomere heterogeneity, or telomere fusions (Supplementary Figure 8). However, we found a variable but elevated incidence of telomere doublets, or fragile telomeres, which are indicative of telomere replication defects, in cells overexpressing TIN2M and the mutant isoforms TIN2S-K280E, TIN2M-K280E, and TIN2L-K280E (Supplementary Figure 8A). These results support the conclusion that the telomere elongation is due to an effect of TIN2 on TPP1/POT1 function and further suggest that TIN2 participates with TPP1 and POT1 in facilitating telomere replication as well as stimulating telomerase processivity.

## Discussion

We have identified a new isoform of TIN2, TIN2M, and have shown that each of the three TIN2 isoforms cooperate with TPP1/POT1 to stimulate telomerase processivity. We found that the TIN2 isoforms play different roles in telomere length regulation in cells. Our data suggest that TIN2 forms a functional shelterin subcomplex with TPP1/ POT1. Considering TIN2 as part of the telomerase processivity complex provides a new way to think about its role in telomere length regulation.

The mutations in *TINF2* in short telomere syndrome patients mostly cluster in a TIN2 domain of unknown function in exon 6 near the C-terminus of TIN2(34, 35). Genetic evidence strongly supports a dominant negative mechanism for the mutant TIN2 proteins. First, TIN2 mutations have autosomal dominant inheritance. The mutant proteins are stably expressed and cause telomere shortening despite the presence of a wild-type TIN2. Secondly, the clustering of disease associated alleles rather than distribution across the coding sequence suggests these are not simply inactivating mutations but rather a gain of function. Finally, there is evidence for selection against the mutant proteins in the hematopoietic lineage *in vivo*(37). The dominant negative mechanism is also supported by experimental evidence(45), but the molecular nature of this effect is not well understood.

### TIN2 C-terminus plays an essential role in telomere length regulation

We found that all three TIN2 isoforms form a complex with TPP1/POT1, stimulate telomerase processivity, and localize to telomeres, yet have different effects on telomere length in cells, underscoring the important role of the TIN2 C-terminus. TIN2 can be divided into two regions: the shelterin-interacting region in the N-terminus, and the C-terminal region that includes the patient mutation cluster, the variable C-terminal extensions of TIN2M and TIN2L, and several other interaction sites and modifications(47, 49, 57, 58) (Figure 1B). All three isoforms contain the shelterin-interacting domain and patient mutation cluster, varying only in the C-terminal extension after E354.

The structure is known for much of the shelterin-interacting region, including the N-terminal TRF2/TPP1 binding domain (TIN2_1-202_)(59) and the short TRF1-interacting motif (TIN2_256-276_)(60). There is no structural information, however, for the C-terminal region, including both the mutation hotspot and the variable C-terminal extension, which contains a highly conserved region with a CK2 phosphorylation site at S396(49). Interestingly, some of the patient mutations are truncations, such as K280X, that generate a short stable protein missing the entire C-terminal region(39). Previous work indicated the importance of the TIN2 C-terminal region, including the high degree of conservation the variable C-terminal extension and the dominant effects of C-terminal truncating mutations. Our work further supports this idea, with discovery of a TIN2 isoform with an alternative C-terminal region and evidence that overexpression of only the isoforms containing C-terminal extensions strongly affect telomere length (Figure 4C,D and (49)). The TIN2 C-terminus may function through binding a novel partner, or through a conformational or structural role.

### TIN2 cooperates with TPP1/POT1 to stimulate telomerase processivity

Our studies indicate that TIN2 is part of the telomerase processivity factor (Figure 5A). All three TIN2 isoforms formed a stable complex with TPP1/POT1 and TERT and further stimulated telomerase processivity over that of TPP1/POT1 alone. This stimulation of processivity required TPP1 and POT1, as there was no stimulation in cells expressing TPP1 TEL-patch mutants or TERT alone (Figure 3 and Supplementary Figure 6). TIN2 could enhance telomerase processivity by improving the TPP1/POT1 complex stability or its interaction with telomerase, or by promoting the telomeric ssDNA interaction of the complex, or some combination of these (Figure 5A).

Interestingly, the identification of TIN2 as an additional component to an already known processivity factor is reminiscent of recent findings in *Tetrahymena*. The *Tetrahymena* telomerase holoenzyme structure(61) revealed previously unknown subunits, Teb2 and Teb3, that interact with the previously defined Teb1-p50 processivity complex. The addition of these proteins to *in vitro* reactions further stimulated telomerase processivity, possibly by stabilizing the complete, assembled, processive enzyme complex(62). Further, this structure revealed that the telomerase holoenzyme contains two single-stranded DNA binding complexes: the p50/TEB processivity factor, which stimulates telomeric G-strand synthesis by telomerase, and the CST complex, which stimulates telomeric C-strand synthesis by lagging strand replication machinery. This is the first evidence of physical coupling of two telomere maintenance processes that have long been known to be coupled *in vivo* (reviewed in(63)).

Our results with TIN2 parallel the discovery of the missing components of the TEB processivity complex in *Tetrahymena*(61, 62), suggesting that TIN2 binding to TPP1/POT1 stabilizes the complex and thus promotes processivity. Interestingly, CST (CTC1/STN1/TEN1), a second ssDNA telomeric complex, interacts with TPP1/POT1 to limit telomere extension by coupling C-strand to G-strand synthesis(64, 65). The C-terminal region of TIN2 is a candidate for coupling TIN2/TPP1/POT1 with CST for coordinated C- and G-strand synthesis, affecting both positive and negative telomere length regulation. Decreased telomerase activity leads to gradual telomere shortening over many generations. Partial uncoupling of telomerase elongation from C-strand synthesis, however, could cause unrestrained telomerase elongation of telomeres, while complete uncoupling could result in telomere shortening by failure to synthesize either C- or G-strands.

### TIN2/TPP1/POT1 is a telomere specific single-stranded binding complex involved in telomere extension and replication

Our data, in combination with previously published work, suggest that TIN2/TPP1/POT1 is a shelterin subcomplex. TIN2 not only increases the processivity stimulation of the complex but also promotes its telomeric localization *in vivo* (Figure 5A). Evidence from previous work supports this conclusion. First, when TRF1 is removed from telomeres by tankyrase-1 modification, TIN2/TPP1 remain at telomeres (13). Second, posttranslational depletion of TIN2 by Siah2 ubiquitination removes TPP1 but not TRF1 or TRF2 from telomeres(57). Further evidence in TIN2 floxed mouse cell lines or TIN2 knockdown in HeLa cells show reduced telomeric TPP1/POT1 localization(26, 27). Similarly, disruption of the TIN2 TRF1-binding motif does not disrupt TRF1, TRF2, or Rap1 localization, but prevents TIN2/TPP1/POT1 accumulation at telomeres in mouse cells(25). Deletion of the TPP1-binding region from mouse TIN2 also prevents localization of TPP1/POT1 to telomeres(28). Finally, genetic evidence using CRISPR knockouts in human cells led the authors to conclude that TIN2/TPP1/POT1 is a shelterin subcomplex(30). These findings together with our work showing the stimulation of telomerase processivity, further supports the conclusion that TIN2, TPP1, and POT1 function together as a subcomplex of shelterin.

The TIN2/TPP1/POT1 heterotrimer likely affects both telomerase and replication fork progression. Considering TIN2/TPP1/POT1 as a telomere specific ssDNA binding (SSB) protein complex helps explain defects in telomere replication that have been reported for both POT1 and TPP1 knockdowns and mutants(12, 13, 31, 44, 56). While most diagrams draw TPP1/POT1 bound to the G-strand overhang at telomeres, this telomere specific SSB complex can also bind the telomeric G-strand exposed during DNA replication(26, 66) (Figure 5B).

TPP1 and POT1 have both been reported to facilitate DNA replication through telomeric tracts(64, 67–69). POT1 mutants that cannot bind DNA cause telomere replication fork stalling, fragile telomeres, and ATR activation(69), possibly due to ssDNA exposure at the telomeric replication fork. TIN2 knockdown(26) and mouse mutants(46) also cause an ATR-mediated DNA damage response. We found that overexpression of some of the TIN2 isoforms resulted in fragile telomeres indicative of telomere replication defects (Figure S8). Taken together, this suggests that the telomeric TIN2/TPP1/POT1and CST complexes may participate directly in replication fork progression through the telomere, and that perturbation of this function may lead to replication fork collapse and activation of ATR.

The interpretation of TIN2/TPP1/POT1 as a ssDNA binding shelterin subcomplex provides an updated view of TIN2’s role in telomere length regulation. We found that TIN2 is expressed as multiple isoforms that have different effects on telomere length in human cells. Strikingly, we found that TIN2 is a previously unappreciated component of the telomerase processivity complex. All three isoforms stimulated telomerase processivity in a TPP1/POT1 dependent manner. Further biochemical work on this heterotrimeric ssDNA telomere binding protein will elucidate the mechanism of TIN2 regulation of telomere length and how it is disrupted in short telomere syndromes.

## Acknowledgements

We would like to thank Drs. Jonathan Alder, Deborah Wuttke, Sarah Wheelan and Leslie Glustrom for suggestions and help with experiments; Dr. Mary Armanios for human LCL cell lines; and Dr. Andrew Holland for FLP-in cell lines. We thank Jonathan Alder, Valerie Gaysinskaya, and Deborah Wuttke for critical reading of the manuscript. This work was supported by NIH Grants R37AG009383 and R35CA209974 and to C.W.G. and a Turock Scholar award to A.M.P.

## Author Contributions

A.M.P and C.W.G designed the project and wrote the manuscript. A.M.P. performed all cloning, cell line generation, telomerase assays, and data analysis. M.A.S. performed immunofluorescence and q-FISH and passaged cell lines. J.P.O and A.M.P. performed 3’RACE and PacBio sequencing. C.J.C. prepared genomic DNA and performed Southern Blots.

## Materials and Methods

### Cell Culture

Cell lines were cultured in the indicated media supplemented with 10% heat-inactivated FBS (Invitrogen, 16140071) and 1% penicillin/streptomycin/glutamine (PSG, Invitrogen 10378016). HeLa, HeLa TREx FLP-in, 293T, and 293TREx FLP-in cells were cultured in DMEM (Gibco); hTERT-RPE1 cells were cultured in DMEM/F12 (Corning); lymphoblastoid cell lines (LCLs) derived from healthy controls (samples obtained after written informed consent and approval from Johns Hopkins Medicine Institutional Review Board) were cultured in RPMI (Gibco); and K562 cell lines were cultured in IMDM (Gibco).

### Expression Constructs

TIN2S cDNA was purchased from Invitrogen (Ultimate ORF IOH80607) in pENTR221. A synthetic gBlock (IDT) containing the downstream TIN2 sequence was used in Gibson Assembly to generate TIN2L. TIN2M was cloned from RT-PCR of endogenous transcripts. TIN2S, M, and L were amplified with primers containing HindIII and NotI restriction sites and an N-terminal myc tag and cloned into pcDNA5/FRT. *TINF2*, the TIN2 full-length gene inclusive of introns, was cloned into pcDNA5/FRT as described in(37). Patient mutations were generated by Site-Directed Mutagenesis. All constructs and mutants were sequence verified by Sanger sequencing at the JHU Synthesis & Sequencing Facility.

P3x-Flag-POT1-cDNA6/Myc-HisC, p3x-Flag-TPP1_87-544_-cDNA6/Myc-HisC, p3x-Flag-TERT-cDNA6/Myc-HisC were a kind gift from the Cech lab(20). We introduced E169A/E171A mutations with site-directed mutagenesis to create TPP1^TEL^. TPP1 or TPP1^TEL^, POT1, and TERT were assembled into a single expression cassette connected by 2A peptides (Supplementary Figure 3) TERT alone was also cloned into pcDNA5. The 2A peptides leave a small tag on the downstream proteins, so TERT was cloned in the last position because it is nonfunctional with C-terminal tags(70–72). Expression cassettes are flanked by BstBI and NotI restriction sites.

### Multiple Sequence Alignments

TIN2 sequences from vertebrates with known or predicted TIN2 proteins were obtained from NCBI. The longer isoform was chosen for organisms with multiple reported isoforms. Sequences were uploaded to PRALINE multiple sequence alignment using the default parameters(73, 74). To make the sequence conservation heat map, PRALINE output was imported into Microsoft Excel, and the alignment scores (0-10) of human TIN2 were colored from white=0, not conserved to navy=8-10, highly conserved.

Sequences used are listed in Supplementary Table 1.

### CRISPR editing

Guide RNAs were selected using the Zhang Lab CRISPR design tool (http://crispr.mit.edu/). For endogenous tagging of TIN2, the guide cgccaccaggggcgtagccaTGG was cloned into pX459-U6-Chimeric_BB-CBh-hSpCas9-2A-Puro. The repair template was generated by PCR from the cloned myc-*TINF2* construct (Supplementary Figure 1). 1μg of Cas9-2A-Puro+TIN2 guide was transfected into 293T cells with 10 molar equivalents of the repair template using XtremeGENE9 (Roche, 6365787001). Editing was enriched with puromycin, cloned by limiting dilution, and screened by PCR and restriction digest. Positive clones were examined by western blot. While we found many edited clones, 293T cells are hypotriploid with an unstable karyotype, and we observed high endogenous Myc expression that interfered with western blotting for myc-tagged TIN2 (Supplementary Figure 1). These caveats make it difficult to further study TIN2 in these knock-in cell lines.

### 3’RACE and PacBio

The 3’RACE and sequencing was performed using samples from five human cell lines (293T, HeLa, RPE-1, K562, LCL) and two mouse samples (CAST/EiJ MEFs, C57BL/6 liver). All mouse samples were obtained under approval by the Institutional Animal Care and Use Committee at the Johns Hopkins University School of Medicine. We combined 3’RACE with Pacific Biosciences (PacBio) Single-Molecule, Real-Time (SMRT) sequencing to cover transcripts from the 5’UTR through the polyA tail. First, we isolated mRNA >10^6^ cells using the RNeasy Kit (Qiagen, 74104) per manufacturer instructions, QIAshredder spin columns (Qiagen, 79654), on-column DNase digestion (Qiagen, 79254) to remove any genomic DNA, and an RNA clean-up. Then we reverse transcribed 1.5 μg mRNA with an oligo-dT_20_ primer with an adapter sequence (GACTCGAGTCGACATCG-T_20_) using the SuperScript III First Strand Synthesis Kit (Qiagen, 18080-051). 5 μl of the resulting cDNA was amplified with Hot Start Phusion Polymerase (Thermo, F-549L) using primers to the adapter and the 5’-UTR (CGGCGACGTTTAAAGCTGA). 3-5 replicate PCR reactions were combined, purified with the QIAquick PCR Purification Kit (QIAGEN, 28104), and submitted to the Johns Hopkins Deep Sequencing & Microarray Core Facility for sequencing. Quality control was performed on a 1:200 dilution of samples using a Bioanalyzer (Agilent, G2939A) High Sensitivity DNA Assay. Products were size selected for the expected size range of 1-3kb. 1 SMRT cell was sequenced per sample. Sequencing reads were processed in the SMRT Analysis v4.0 software, aligned to chromosome 14 with HISAT2 and assembled into potential transcripts using StringTie(75, 76). StringTie was first run for individual samples using the default settings except the minimum isoform fraction was set to 0.01 instead of 0.1. To build a gene model for all human reads, StringTie --merge was run with the minimum isoform fraction set to 0.05. HISAT2 and StringTie results were viewed in IGV(77, 78).

### Western Blotting

Cells were lysed on ice in CHAPS lysis buffer (10 mM Tris-HCl, 1 mM MgCl_2_, 1 mM EGTA, pH 8.0, 0.1 mM Benzamidine, 5 mM β-Mercaptoethanol (BME), 0.5% CHAPS, 10% Glycerol, pH 7.5) and clarified by centrifugation. Samples were denatured with 1X LDS (Invitrogen, NP0008) with 50 mM DTT and heated at 65°C for 10 minutes and separated on a 4-12% Bis-Tris gel (NuPAGE, NP0323) in 1X MOPS buffer (Invitrogen, NP0001) with 3 μl of SeeBlue Plus2 (Thermo, LC5925) prestained ladder to estimate molecular weight. Proteins were transferred to PVDF, blocked in 1X TBS + 0.1% Tween20 (TBST), 5% milk (Bio-Rad 170-6404), probed with the indicated antibodies, and developed by chemiluminesence with an ImageQuant LAS4000 imager (GE Healthcare). Primary antibodies and concentrations are as follows: mouse anti-myc 4A6 (Millipore 05-24), 1:2000; mouse anti-FLAG M2 (Sigma F1804), 1:5000; rabbit anti-tubulin (Abcam, ab6046), 1:5000. Secondary antibodies were anti-mouse IgG or anti-rabbit IgG conjugated to HRP (Cell Signaling), 1:10,000.

### Co-Immunoprecipitation

Immunoprecipitations were carried out using either anti-c-myc agarose (Pierce 20168) or anti-FLAG M2 affinity gel (Sigma A2220). 20 μl of bead slurry per reaction was washed with PBS and equilibrated in CHAPS buffer before adding 45 μl lysate. Samples were incubated in an end-over-end mixer at 4°C for two hours. Beads were pelleted, washed 4 times with 300 μl 1X CHAPS buffer, and resuspended in 2X LDS loading dye for western blot analysis.

### Telomerase Assays

Telomerase assay cell lines were generated in 293 TREx FLP-in cells (Invitrogen, R78007) as described(79). Briefly, parental cells were transduced with a telomerase RNA (TR) lentivirus, selected, and cloned by limiting dilution. Then, the TPP1/POT1/TERT or TPP1^TEL^/POT1/TERT construct was integrated at a single site in a TR overexpressing clone using the Flp-in system (Invitrogen). For telomerase assays, 5 × 10^5^ cells of the respective cell line were plated in each well of a 6-well dish. The next day, the indicated 2.5 μg of the indicated TIN2 or GFP construct was transfected with Lipofectamine 2000 (Invitrogen, 11668019) following the manufacturer’s protocol. After 48 hours, cells were lysed in 100μl 1X CHAPS lysis buffer and clarified by centrifugation. Telomerase assays were performed as described in(79) using 5 μl of clarified cell lysate. Assays were quantitated in ImageQuantTL (GE Healthcare) using the 15+ method as described(20). Statistical analysis was performed in GraphPad Prism.

### TIN2 Overexpression Cell Lines

HeLa FLP-in cells were seeded in 6-well plates with 3 wells per construct. The next day, each well was transfected with 100 ng pcDNA5/FRT-TIN2 or -GFP construct and 900 ng pOG44 (FLP-recombinase). Wells were then pooled and selected for integration of the pcDNA5/FRT plasmid with hygromycin for 14 days. After selection, isogenic clones (> 20 per cell line) were pooled and released from selection (“week 0”). Cells were split 1:10 three times a week. No growth difference was detected. Parental HeLa FLP-in cell line was validated with STR profiling through the Johns Hopkins Genetic Resources Core Facility.

### Immunofluoresence

HeLa FLP-in cells were plated in chamber slides. The following day, media was removed, the cells were washed with PBS and fixed with 4% paraformaldehyde (PFA) for 20 minutes. Slides were washed with PBS, treated with 0.5% Triton in PBS for 15 minutes, washed with PBS and blocked in 10% goat serum in PBS for 30 minutes (Sigma, G0923). Slides were incubated with a mixture of both primary antibodies for 1 hour at room temperature, washed with PBS and incubated with a mixture of both secondary antibodies for one hour at room temperature. After washing with PBS coverslips were mounted with DAPI/Vectashield. Antibodies and dilutions are as follows: mouse anti-myc clone 4A6 (Sigma, 05-724) 1:200, rabbit anti-TRF2 (Novus Biologicals, NB110-57130) 1:800, goat anti-mouse IgG1-AlexaFluor 488 (Invitrogen, A21121) 1:400, and goat anti-rabbit IgG-AlexaFluor 555 (Invitrogen, A21429) 1:400. Slides were imaged on a Nikon Eclipse Ni-E microscope with a 60x objective using the NIS Elements software.

### Telomere Southern Blots

Genomic DNA was prepared from ~3-6 × 10^6^ frozen cell pellets lysed in Nuclei Lysis Solution (Promega, A7941), treated with RNAse A (10mg/ml, Roche) and overnight with Proteinase K (400mg/ml, ThermoFisher), followed by salting out of the proteins with Protein Precipitation Solution (Promega, A7951). The genomic DNA was precipitated with isopropanol and resuspended in TE (10mM Tris pH8.0, 1mM EDTA). Approximately 2μg of genomic DNA, quantitated by a Qubit 3.0 Fluorometer (Life Technologies), was cut with the restriction enzyme with MseI (NEB, R0525M) overnight, run on a Southern blot, and hybridized with a radiolabeled telomere fragment from JHU821 as described(79). Images were captured, converted, and quantitated from Storage Phosphor Screens (GE Healthcare) as described in(79).

### Telomere analysis by q-FISH

For metaphase Fluorescent in situ hybridization (FISH) analysis, cultures were first arrested in with Karyomax Colcemid (Invitrogen) for 6-7 h. The cells were trypsinized in 0.05% Trypsin-EDTA (Gibco), washed in PBS, swelled with 0.075M KCl at 37°C for 15 min and fixed in methanol:acetic acid (3:1). Cell suspensions were then dropped onto chilled slides and dried overnight. FISH was performed using a Cy3-labeled (CCCTAA)_3_ PNA oligonucleotide (PE Biosystems). Metaphase spreads were counterstained with DAPI/Vectashield. Slides were blinded during image acquisition and analysis. Images were acquired using a Nikon Eclipse NI-E microscope and NIS Elements software. Telomere fusions, fragile telomeres, and signal-free ends were tallied in 10-metaphases per sample in a total of three replicates. Telomere length was measured in TFL-Telo V2.0, and outliers were analyzed as described(80) to determine PQ ratios and sister-telomere ratios. Histograms of telomere lengths were generated in GraphPad Prism.

